# *Plasmodium falciparum* Niemann-Pick Type C1-Related Protein is a Druggable Target Required for Parasite Membrane Homeostasis

**DOI:** 10.1101/371484

**Authors:** Eva S. Istvan, Sudipta Das, Suyash Bhatnagar, Josh R. Beck, Edward Owen, Manuel Llinás, Suresh M. Ganesan, Jacquin C. Niles, Elizabeth A. Winzeler, Akhil B. Vaidya, Daniel E. Goldberg

**Author notes:** Co-first authors (Eva S. Istvan, Sudipta Das).

## Abstract

Plasmodium parasites possess a protein with homology to Niemann-Pick Type C1 proteins (*Plasmodium falciparum* Niemann-Pick Type C1-Related protein, PfNCR1). We isolated parasites with resistance-conferring mutations in PfNCR1 during selections with three diverse small-molecule antimalarial compounds and show that the mutations are causative for compound resistance. PfNCR1 protein knockdown results in severely attenuated growth and confers hypersensitivity to the compounds. Compound treatment or protein knockdown leads to increased sensitivity of the parasite plasma membrane (PPM) to the amphipathic glycoside saponin and engenders digestive vacuoles (DVs) that are small and malformed. Immuno-electron microscopy and split-GFP experiments localize PfNCR1 to the PPM. Our experiments show that PfNCR1 activity is critically important for the composition of the PPM and is required for DV biogenesis, suggesting PfNCR1 as a novel antimalarial drug target.

## Introduction

Several whole-parasite chemical library screens have identified thousands of compounds with potent antimalarial activity^1,2^. To facilitate drug development, it is important to identify targets of these compounds. Target identification can be extremely challenging, especially in organisms like *Plasmodium* that contain large numbers of proteins with unknown function. Evolution of compound-resistant malaria parasites can be helpful in the discovery of the molecular mechanisms by which compounds kill the organism^3–6^.

In this study, we investigated a gene that acquired single nucleotide polymorphisms (SNPs) or was amplified in selections with three diverse compounds. PF3D7_0107500 encodes a membrane protein with sequence motifs found in Niemann-Pick C1 (NPC1) proteins. Human NPC1 (hNPC1) has been the subject of numerous studies because of the protein’s importance in cholesterol egress from late endosomes^7^. Patients with mutations in hNPC1 suffer a fatal neurodegenerative lipid storage disorder characterized by the accumulation of lysosomal cholesterol, sphingomyelin, as well as other lipids^8^. Niemann-Pick C1-Related (NCR1) proteins are conserved in eukaryotic evolution and are most easily identified by their membrane domains^9^. In humans, NPC1 accepts cholesterol from its partner protein, the high affinity cholesterol-binding protein NPC2^10^. NCR1 homologs are also present in organisms that do not contain readily identifiable NPC2 proteins or internalize sterol by endocytosis. Based on studies with yeast NCR1, Munkacsi et al. proposed that the primordial function of NCR1 is the regulated transport of lipophilic substrates such as sphingolipids^11^.

Until now the function of PF3D7_0107500, which we call *Plasmodium falciparum* Niemann-Pick Type C1-Related protein (PfNCR1), has been unclear. In this study, we prepared a genetic knockdown (K/D) of *pfncr1* and showed that K/D critically slows blood-stage parasite replication. Furthermore, *pfncr1* K/D caused parasites to become abnormally sensitive to the pore-forming amphipathic glycoside saponin. Treatment with any of the three compounds that we identified during resistance selection phenocopied the gene knockdown, suggesting that the compounds interfere with PfNCR1 function. Here we show that PfNCR1 is druggable and necessary for maintaining the proper membrane lipid composition of blood-stage parasites.

## Results

### Mutations in PfNCR1 Provide Resistance to Three Diverse Compounds

As part of a study aimed at analyzing the *P. falciparum* resistome, we isolated parasites resistant to three structurally diverse compounds (Fig. 1A) that contained mutations in one common gene, PF3D7_0107500^12,13^, which is predicted to encode a 1470 amino acid membrane protein. Sequence similarity searches indicated homology to a protein previously studied in the related apicomplexan parasite *Toxoplasma gondii* called Niemann-Pick Type C1-Related Protein (TgNCR1). Lige et al. identified sequence elements conserved between TgNCR1 and hNPC1, a lysosomal integral membrane protein^14^. The same sequence elements are also present in PfNCR1. Cryo-EM and crystal structures of hNPC1 reveal a 13-helix transmembrane region containing a sterol-sensing domain (SSD) (orange) and a conserved C-terminal transmembrane domain (C-TM) (magenta) (Fig. 1B)^8,10^. The C-terminal targeting sequence that extends past the C-TM in hNCR1 and localizes this protein to the lysosome, is not present in PfNCR1. Lumen-exposed domains (grey and blue in Fig. 1B) complete the hNPC1 structure. Sequence similarity between hNPC1 and PfNCR1 is restricted to portions of the transmembrane region (orange and magenta) and to approximately 45 amino acids N-terminal to the SSD (red). Based on this limited sequence similarity, we generated a cartoon model of PfNCR1 (Fig. 1C). We observed five mutations in our compound-resistant parasites: A1108T came from selections with MMV009108; M398I and A1208E from selections with MMV028038; and S490L and F1436I from selections with MMV019662. The model suggests that three of the mutations are proximal to the membrane domain, while the other two localize to the hydrophilic domains. We used single-crossover allelic exchange to introduce one mutation from each resistance selection into a clean genetic background (Supplementary Figs. S1A, S1B, S1I). With this strategy, PfNCR1 is expressed from its native promoter and contains a C-terminal green-fluorescent protein (GFP) tag in addition to the mutation. We also generated non-mutated allelic exchange control parasites containing the GFP tag. Inclusion of the C-terminal GFP did not alter the sensitivity to MMV009108 (Fig. 1D), while parasites with single mutations in PfNCR1 were resistant to the compounds with which they were selected (Fig. 1D-F).

**Figure 1:**
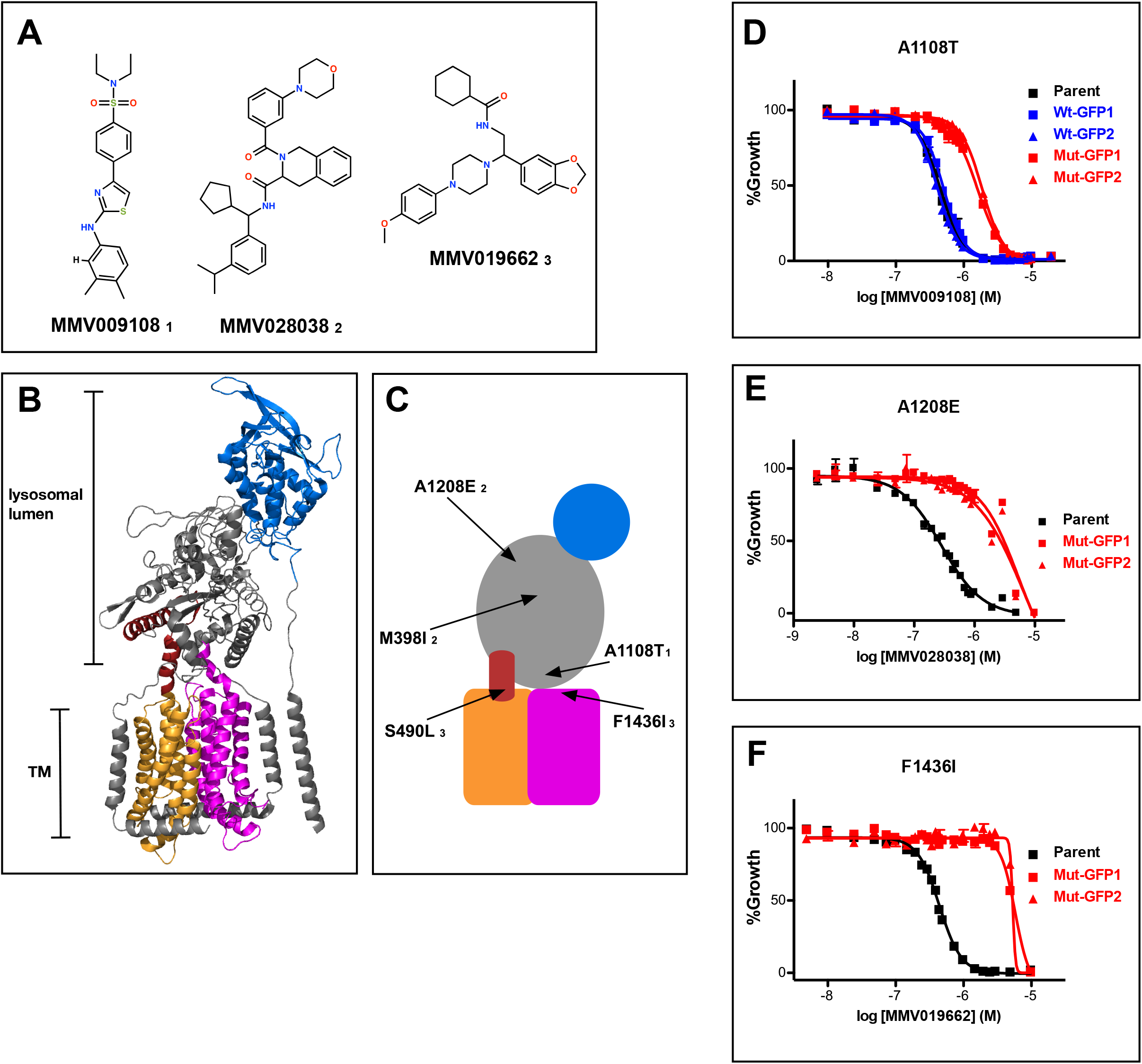
Mutations in PfNCR1 confer resistance mutations to three antimalarial compounds. A) Structures of the three structurally diverse compounds that yielded resistant mutations in PfNCR1. The lower-case numbers next the compound IDs are used in C) to match mutations with specific compounds. B) Ribbon model of the structure of hNCR1 solved by cryoEM^8^. PDB coordinates: 3JD8. The SSD is shown in orange, the conserved C-terminal membrane domain is shown in magenta, the domain that interacts with hNCR2 is in blue and an additional sequence stretch with similarity to PfNCR1 is in red. C) Cartoon model of the possible domain arrangement in PfNCR1. Sequence similarity to hNCR1 is restricted to the red, orange and magenta domains. Locations of resistance-conferring mutations are shown with arrows. Compound IDs matching with mutations are shown in lower case numbers and match Fig. 1A. D-F) Concentration response curves of blood-stage parasites (all in 0.1% DMSO) measured using a flow cytometry-based assay. Each panel shows a different compound and a different mutation. D) MMV009108, E) MMV02803, F) MMV019662. Black = parental 3D7 parasites; red = two independent clones of parasites with mutant allelic exchange; blue = two independent clones of parasites with wild-type allelic exchange (for part D only). The error bars (S.D.) for a representative experiment (biological triplicates) are shown and are very small. This experiment was done two times.

We examined the effect of single mutations on the different compounds. A1108T or F1436I mutant parasites were resistant to all three compounds (Figs. 1D and 1F, Supplemental Figs. S1C-F), while A1208E mutant parasites were sensitive to the two compounds that were not used in the A1208E selection (Supplemental Figs. S1G, S1H). These findings suggest that amino acids modeled to be proximal to the membrane domain (A1108 and F1436) may have some functional overlap, while the putative soluble domain A1208 may have a different activity.

### PfNCR1 is Important for Asexual Parasite Viability and is Targeted by Antimalarial Compounds

An attempt to disrupt the *pfncr1* gene using a CRISPR/Cas9-targeting approach did not succeed, suggesting an essential function during blood-stage malaria growth. Next, we created parasites in which *pfncr1* expression is regulated by anhydrotetracyline (aTc) using the previously described TetR-DOZI/aptamer translational repression technology^15,16^ (Supplementary Figs. S2A, S2B). When we removed aTc from highly synchronized, young ring-stage parasites, PfNCR1 expression was reduced within the same cell cycle and undetectable in the following cell cycles, as judged by western blots to detect a C-terminal hemagglutinin (HA) sequence on the aptamer-tagged parasites (Fig. 2A). While protein levels after aTc withdrawal were affected almost immediately, parasite replication rates decreased only after 3-4 days (Fig. 2B, inset). After this slow onset of reduced growth, PfNCR1 knockdown clearly resulted in markedly less fit parasites. Complementing knockdown parasites with a second copy wild-type PfNCR1 rescued the growth defect (Fig. 2C, Supplementary Fig. S2C). Modulating the expression level of PfNCR1 with aTc shifted the MMV009108 concentration-response curve (Fig. 2D) and maximal knockdown hypersensitized parasites to the three compounds that were used for the resistance selection (Fig. 2E-G), while sensitivity to the antimalarial mefloquine was not affected (Fig. 2H). Our findings suggest that PfNCR1 performs a function important for the viability of blood-stage malaria parasites and that the three compounds act directly on the protein.

**Figure 2:**
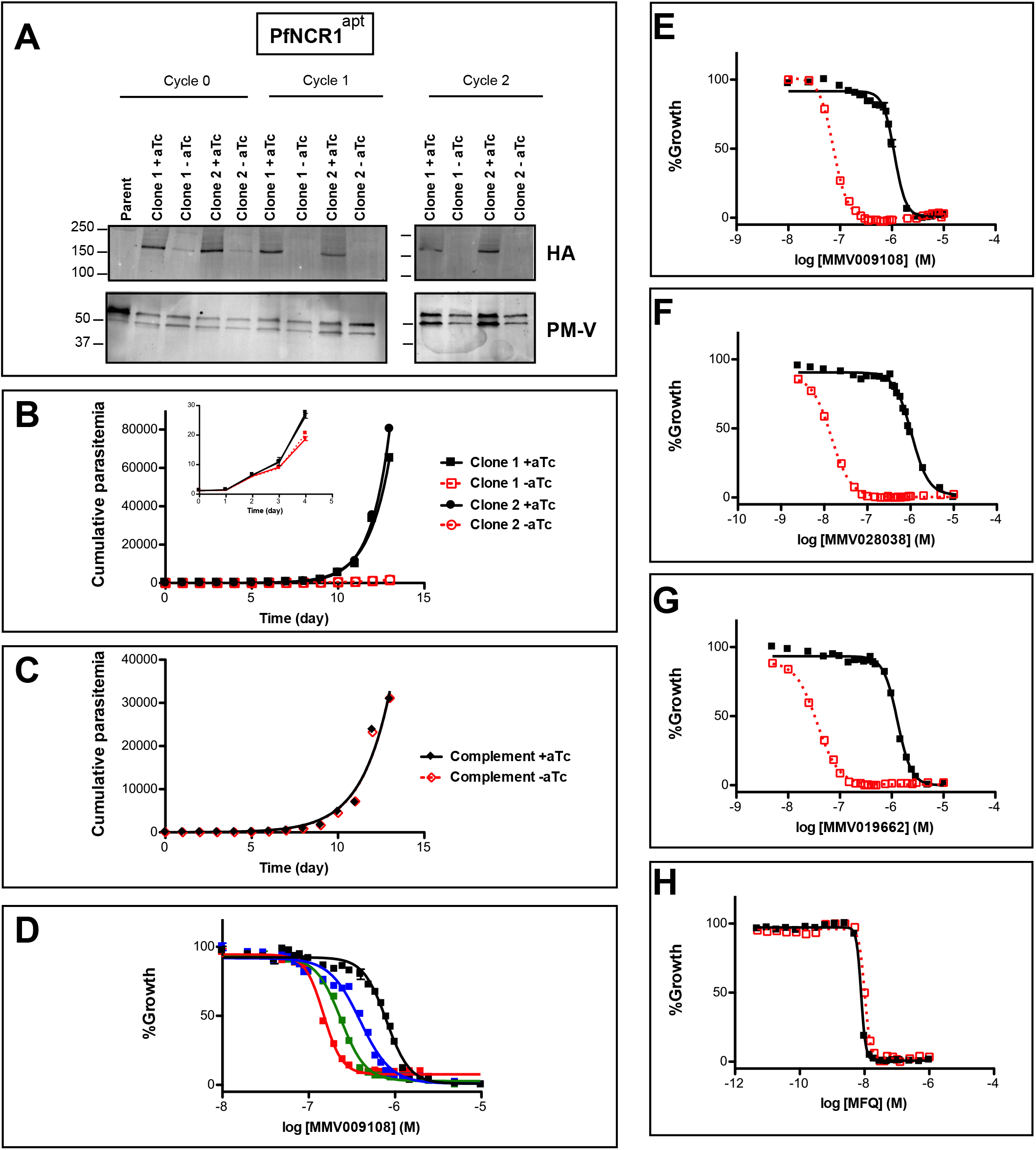
PfNCR1 is required for blood-stage parasite replication and is targeted by three antimalarials. A) Western blot showing regulation of the PfNCR1^apt^ by aTc. Trophozoite-stage parasites were harvested from the replication cycle in which aTc was removed (cycle 0), as well as the following two cycles. PfNCR1 was detected using a C-terminal HA-tag. The ER membrane protein plasmepsin V (PM-V) was used as a loading control. Note that the two bands recognized by α-PM-V antibody correspond to the full-length protein and a proteolytic fragment of the protein produced during the membrane isolation. Expected sizes: 171k Da for PfNCR1-HA, 69k Da for PM-V. This experiment was done two times. B) Replication of PfNCR1^apt^ parasites. Using a flow cytometry assay, the replication of two PfNCR1^apt^ clones was monitored over two weeks. +aTc is in black and solid lines, −aTc is in red and dashed lines. One representative experiment with biological triplicates is shown. The inset magnifies the initial time points. Doubling times in days are as follows (95% confidence intervals in parentheses): clone 1 - aTc=1.596 (1.546-1.650), R^2^=0.9964; clone 1 +aTc=0.8663 (0.8218-0.9159), R^2^=0.9930; clone 2 −aTc=1.463 (1.417-1.512), R^2^=0.9965; clone 2 +aTc=0.7776 (0.7559-0.8005), R^2^=0.9981. This experiment was done four times. C) Complementation of PfNCR1^apt^ rescues growth phenotype. Wild-type PfNCR1 was stably expressed in the PfNCR1^apt^ background. Replication of parasites was monitored over two weeks. +aTc is in black and solid line, −aTc is in red and dashed line. One representative experiment with biological triplicates is shown. Doubling times in days are as follows (95% confidence intervals in parentheses): −aTc=1.152 (1.0.36-1.298), R^2^=0.9658; +aTc=1.166 (1.039-1.329). This experiment was done four times. Note that the complemented strain grows less well than PfNCR1^apt^ with aTc (B), but that there is no significant difference +/− aTc. D) Expression level of PfNCR1 correlates with sensitivity to MMV009108. Concentration response curves using a flow cytometry-based growth assay. After aTc washout, aTc was replenished at different concentrations and parasitemias were measured after 72 hrs. Black=50nM aTc, blue=10nM aTc, green=3nM aTc, red=1nM aTc. Biological triplicates. This experiment was done one time. E-H) PfNCR1 K/D hypersensitizes parasites to three compounds. Concentration-responses of PfNCR1^apt^ parasites to E) MMV009108, F) MMV028038, G) MMV019662, and H) mefloquine (MFQ) (control compound) without aTc (red open symbols, dashed lines) or with 500nM aTc (black symbols, solid lines) after 72 hrs. One representative experiment with biological triplicates is shown. The experiment in E) was done three times the experiments in for F-H) were done two times.

### PfNCR1 localizes to the parasite plasma membrane

To better understand the functional significance of PfNCR1, we localized the protein. For this purpose, we used parasites expressing wild-type PfNCR1 protein tagged with a C-terminal GFP from its native promoter. Live microscopy showed fluorescence surrounding the intraerythrocytic parasites (Fig. 3A). The distribution of GFP was in contrast to an earlier suggestion that PfNCR1 may reside in the digestive vacuole (DV) membrane^17^. Immuno-electron microscopy confirmed localization of PfNCR1 to the membranes surrounding parasites (Fig. 3B, C). Blood-stage parasites are surrounded by two membranes in very close apposition - the parasitophorous vacuolar membrane (PVM) and the PPM. The resolution of our immuno-electron microscopy images was not sufficient to determine whether PfNCR1 is present in the PVM or the PPM. To answer this question, we prepared split-GFP constructs^18,19^ in which GFP strands 1-10 are expressed either in the parasite cytoplasm or targeted to the lumen of the parasitophorous vacuole (PV) and GFP strand 11 is expressed as a C-terminal tag on PfNCR1 (Supplementary Fig. S3). GFP fluorescence was only observed when cytoplasmic GFP 1-10 was co-expressed with PfNCR1-GFP11 (Fig. 3D-G), suggesting that the C-terminal residues of PfNCR1 project into the parasite cytoplasm. Based on these results, we propose a model in which PfNCR1 membrane domains are in the PPM while the soluble domains project into the PV (Fig. 3H). In contrast to hNPC1, PfNCR1 does not appear to localize to internal organellar membranes. Nevertheless, our model suggests that the relative orientation of the cytosolic regions in these two distantly related proteins is conserved.

**Figure 3:**
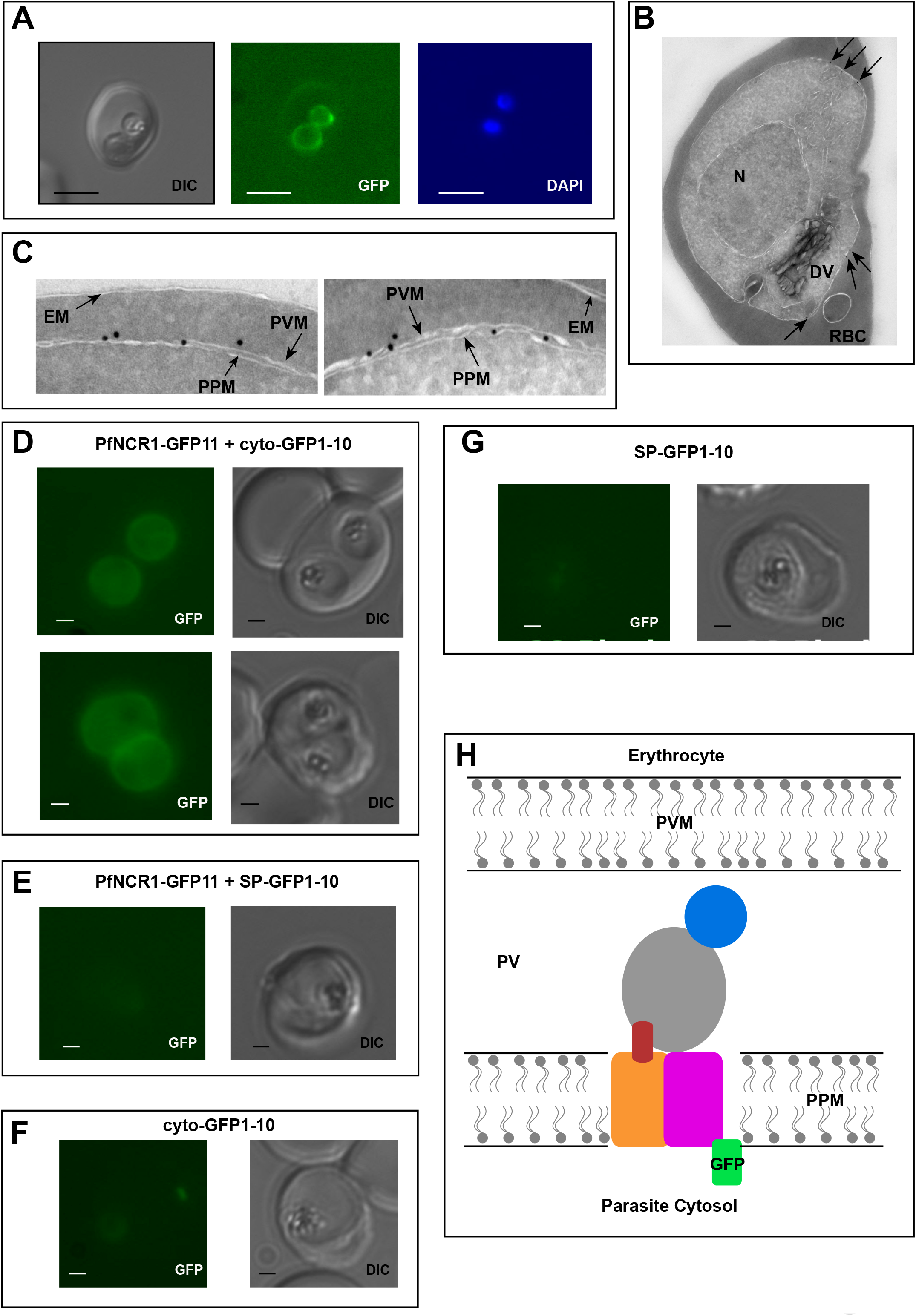
PfNCR1 localizes to the parasite plasma membrane. A) Live fluorescence microscopy with C-terminally GFP-tagged wild-type PfNCR1-expressing parasites (clone Wt-GFP1 from Fig. 1) localizes PfNCR1 to the parasite surface. Scale bar 5μm. B-C) Immuno-electron-micrographs of trophozoite-stage parasites using α-GFP antibody. Arrows mark gold particles, RBC=infected red blood cell, DV=digestive vacuole, N=nucleus. The close-up in C) shows gold particles clustered at the parasite-delimiting membranes. EM= erythrocyte membrane; PVM=parasitophorous vacuolar membrane; PPM=parasite plasma membrane. D-G) Live fluorescence microscopy on split-GFP expressing parasites. D) Co-expression of PfNCR1-GFP11 with cytoplasmic GFP1-10. E) Co-expression of PfNCR1-GFP11 with vacuolar GFP1-10. F) Cytoplasmic GFP1-10 without expression of PfNCR1-GFP11. F) Vacuolar GFP-1-10 without expression of PfNCR1-GFP11. Scale bar: 1μm. H) Cartoon of the proposed orientation of PfNCR1 in the PPM (parasite plasma membrane). PV = parasitophorous vacuole; PVM = parasitophorous vacuolar membrane.

### Compound treatment or protein knockdown hypersensitizes parasites to saponin

Saponins are amphipathic glycosides with high affinity for cholesterol that are capable of penetrating membranes^20,21^. Inhibiting the Na^+^-efflux pump PfATP4 has previously been shown to lead to changes in PPM saponin sensitivity^22^. We were curious whether interfering with PfNCR1 function would have similar effects. We noticed decreased levels of cytosolic aldolase protein in saponin parasite extracts after incubation with MMV009108, while levels of the PVM-localized membrane-bound protein EXP2 did not change (Fig. 4A). Hypersensitivity to saponin was reversed when MMV009108 was removed by washout. We obtained similar results in experiments probing for a different cytosolic protein, haloacid dehalogenase 1 (HAD1) (Supplementary Fig. S4A). Using a flow cytometry-based assay and a previously reported parasite clone expressing eGFP^23^, we observed elevated saponin-induced leakage of cytoplasmic eGFP after incubation of parasites with sub-EC50 concentrations of MMV009108, MMV028038 and MMV019662 (Fig. 4B-D). Western blots probing for eGFP in supernatant and pellet fractions showed that the decrease in signal of cytosolic proteins was not a consequence of increased protein degradation, but rather of elevated leakage of cytoplasmic contents (Supplementary Fig. S4B). These results suggest that the PPM, the membrane to which PfNCR1 localizes, undergoes a redistribution of membrane lipids during compound treatment. We have previously shown that, for PfATP4 inhibitors, induction of saponin sensitivity is abrogated in parasites adapted to grow in low [Na^+^]^22^. This was different from the effect of MMV009108 treatment where we observed saponin hypersensitivity in regular medium, as well as and in low [Na^+^]-containing medium (Fig. 4E). Also, unlike PfATP4 inhibitors, MMV009108 did not result in Na^+^ influx into parasites (Fig. 4F). PfNCR1 K/D was minimally hypersensitive to KAE609 (Supplementary Fig. 5). We conclude that MMV009108 acts directly on PfNCR1 but suggest that PfATP4 activity influences PfNCR1 function (PfATP4 mutants are hypersensitive to our compounds^12^).

**Figure 4:**
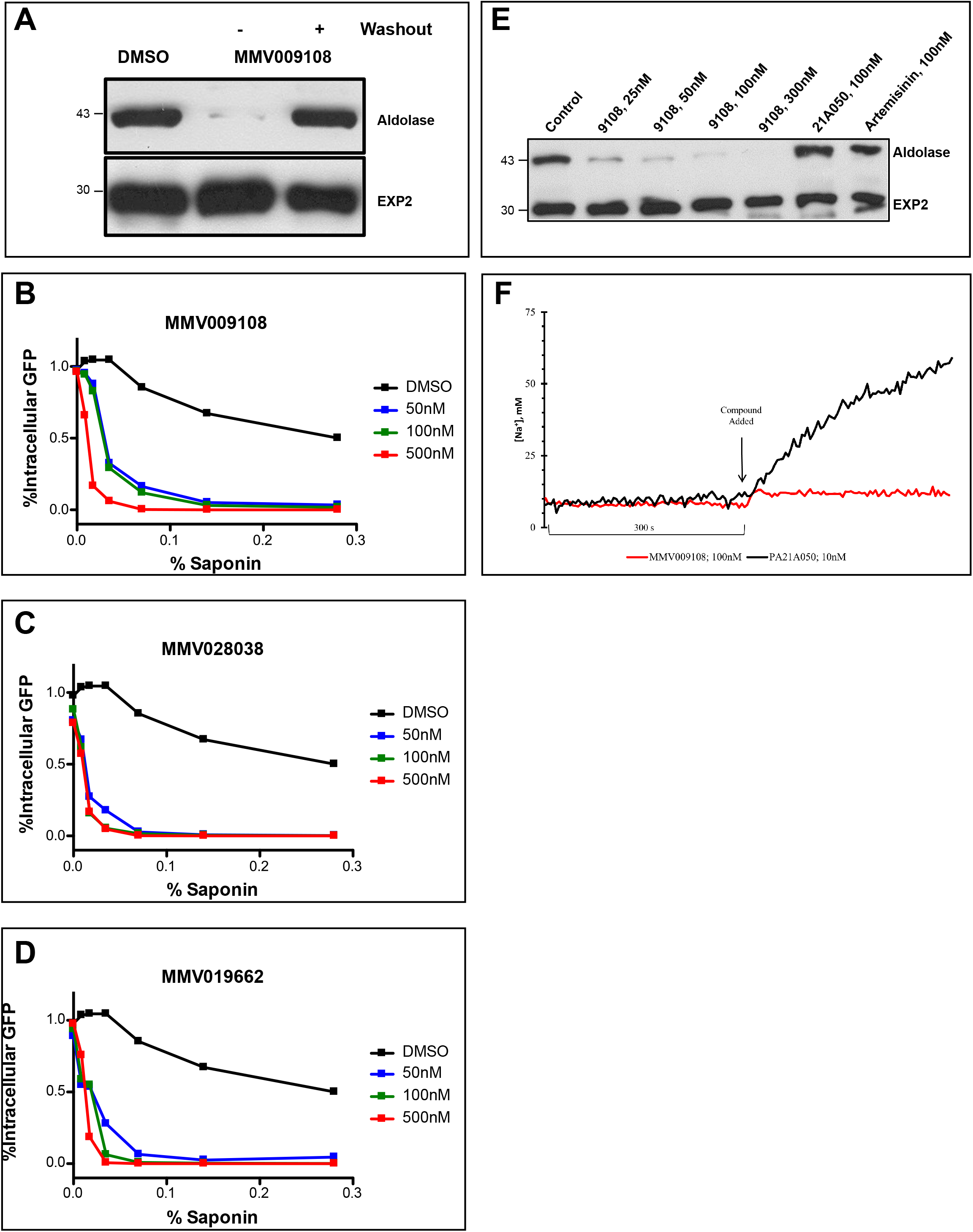
Compound treatment hypersensitizes parasites to saponin. A) Strain 3D7 parasites (30-34h post-infection) were exposed to DMSO or 100nM MMV009108 for 2h. Treated parasites were washed out and rescued by growing in compound free cRPMI medium for another 2h. Parasites were treated with saponin (0.02%) to release parasites followed by western blot analysis using antibodies to parasite aldolase or EXP2. EXP2 was used as a loading control. This experiment was done three times. B – D) Flow cytometry-based assay to monitor cell leakiness using a cytoplasmic GFP expressing parasite clone (NF54^eGFP^). Parasites were incubated with MMV009108 (C), MMV028038 (D), or MMV019662 (E) at the indicated concentrations for 1hr (DMSO was the vehicle control). Following compound washout with PBS, parasites were released from RBCs with saponin. Using flow cytometry, 50,000 cells were counted and scored as GFP positive or negative. At 0% saponin, all samples had similar numbers of GFP-positive cells (~80%). The experiment in C) was done three times. The experiments in D and E) were done two times. E) Low Na^+^-adapted trophozoite stage 3D7 parasites were subjected to varying concentration of MMV009108 for 2 hrs followed by saponin (0.02%) treatment to release the parasites and subjected for western blot analysis using antibodies to parasite aldolase or EXP2 (loading control). 100 nM PA21A050^22^ and 100 nM artemisinin were used as controls. This experiment was done two times. F) Unlike the pyrazoleamide PA21A050, MMV009108 does not induce Na^+^ influx into parasites as judged by SBFI fluorescence ratiometric measurement of intracellular [Na^+^].

We looked for changes in the PVM using a parasite clone in which the fluorescent protein mRuby3 is targeted via a signal peptide to the PV (Fig. 5A). As expected, the PVM was exquisitely sensitive to saponin and mRuby was released irrespective of drug treatment (Fig. 5B). Leakage of cytosolic HAD1 after saponin treatment was enhanced by MMV009108, as previously seen (Supplementary Fig. S6). With the same PV-targeted mRuby parasites we examined the sensitivity of the PVM to the cholesterol-binding toxin tetanolysin, which, at low concentrations, normally lyses the erythrocyte membrane but not the PVM^24^. Treatment with MMV009108 did not alter PVM susceptibility to tetanolysin (Fig. 5C), suggesting that compound treatment does not perturb PVM lipid composition.

**Figure 5:**
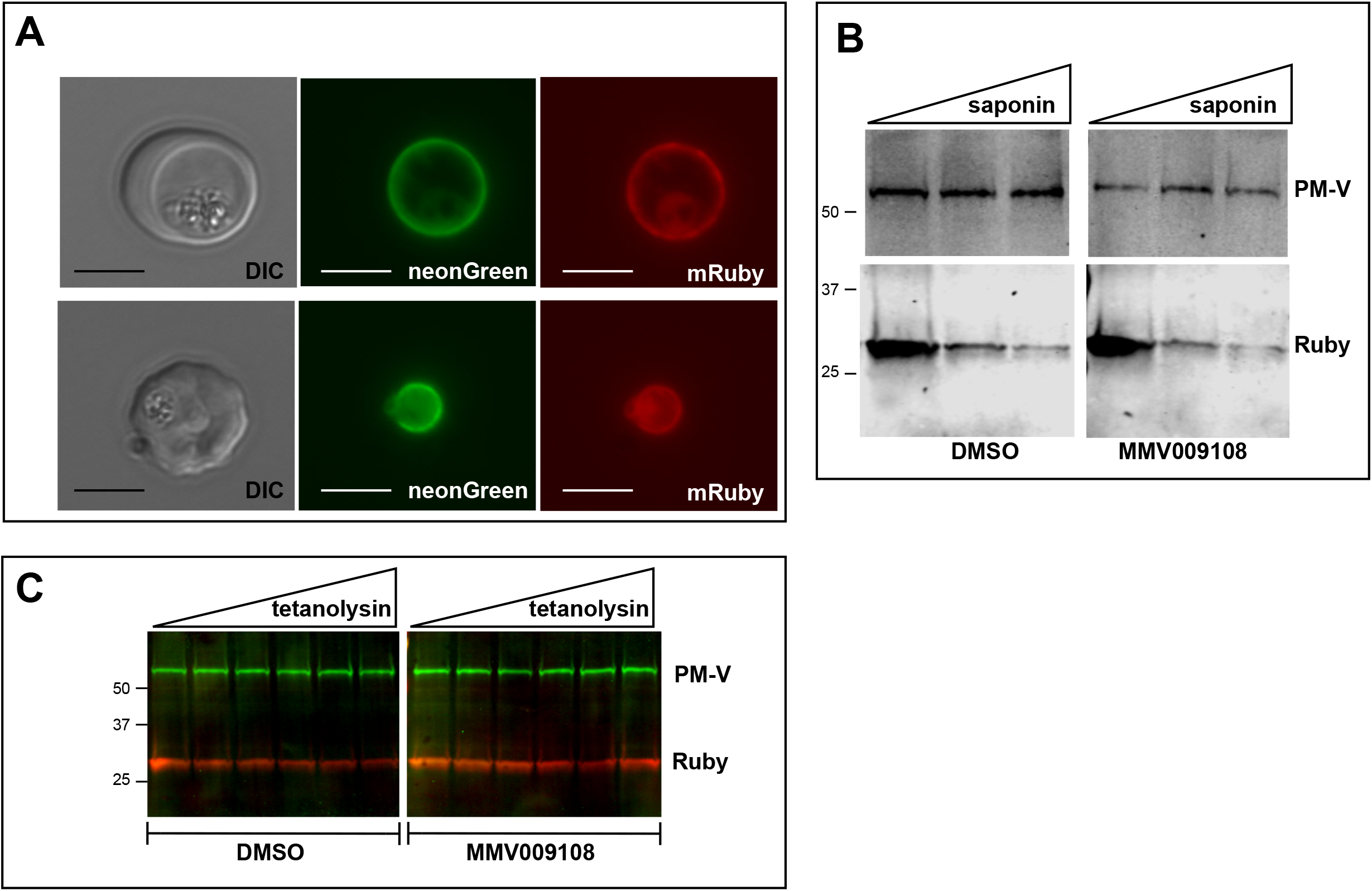
PVM lipid homeostasis is not affected by MMV009108. A) Live microscopy on NF54-EXP2-mNeonGreen+PV-mRuby3 parasites. The PVM protein EXP2 is expressed as mNeonGreen fusion; mRuby3 is targeted to PV lumen. Scale bar=5μm. B) Western blot on saponin-treated NF54-EXP2-mNeonGreen+PV-mRuby3 parasites following treatment with 500nM MMV009108 for 2 hrs. The saponin gradient was as follows: 0%, 0.009%, 0.018%. This experiment was done two times. C) Western blot on NF54-EXP2-mNeonGreen+PV-mRuby3 parasites following treatment with tetanolysin (concentrations: 0, 0.5, 1, 2.5, 5, 7.5ng/ml). Blot was probed with anti-RFP and anti-PM-V antibodies. This experiment was done two times. Expected sizes: PV-Ruby3=27kDa, PM-V=69kDa.

Next, we examined whether PfNCR1^apt^ parasites are hypersensitive to saponin after K/D. Removal of aTc sensitized parasites to saponin as monitored by the loss of cytoplasmic HAD1, while complemented control parasites expressing wild-type PfNCR1 in the knockdown parasite background had normal saponin sensitivity (Fig. 6A). As an independent marker, we prepared a PfNCR1^apt^ parasite line expressing cytosolic eGFP. In this background, PfNCR1 K/D increased PPM sensitivity to saponin within 22 hours of aTc removal (Fig. 6B), much more rapidly than the onset of slowed parasite growth (Fig. 2B). Adding back aTc to PfNCR1^apt^ parasites rapidly restored normal saponin sensitivity (Fig. 6B). Similarly, saponin sensitivity after knockdown of PfNCR1 for 40 hours was reversible in as little as 2 hours (Supplementary Fig. S6). In summary, PfNCR1 K/D phenocopies the effect of the three compounds on the PPM, suggesting that the compounds we identified interfere with PfNCR1 activity and that PfNCR1 function is required to maintain normal PPM lipid composition.

**Figure 6:**
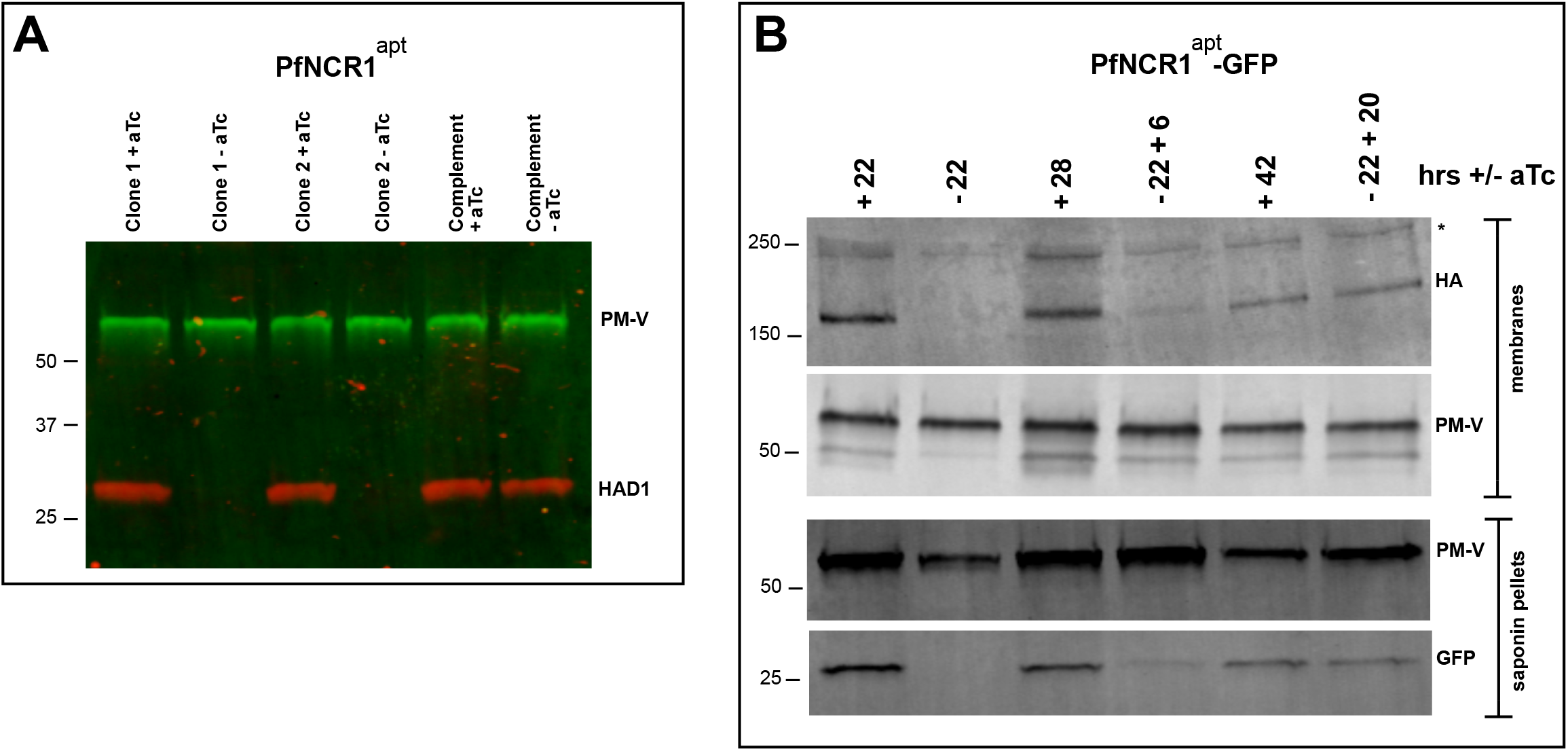
PfNCR1 K/D hypersensitizes parasites to saponin. A) Western blot analysis of saponin extracts (0.07%) from two PfNCR1^apt^ clones and complemented parasites. Parasites were harvested 24 hrs after aTc washout. This experiment was done two times. B) Replenishing aTc after washout reverts the K/D phenotype. aTc was removed from PfNCR1^apt^-GFP parasites (stable expression of cytosolic GFP). 22 hrs after washout, one set of parasites was harvested, while aTc (500nM) was added back to another set of parasite samples for 6 or 20 hrs. Parasites were either harvested to prepare membranes, or released with saponin. Lysates were subjected to Western blotting. * in top blot (anti-HA) marks a crossreacting protein. This experiment was done two times. Expected sizes: HAD1=33kDa, PM-V=69kDa, PfNCR1-HA=171kDa, GFP=27kDa.

### PfNCR1 activity is required for digestive vacuole function

We hypothesized that DV formation could be affected by PfNCR1 impairment as DVs are formed from endocytic vesicles that invaginate at the PPM (Fig. 7A). To observe DVs in live parasites we used a strain that expresses GFP as a fusion protein with the DV protease plasmepsin II (PMII)^25^. In this strain, PMII-GFP is produced as a membrane-bound pro-enzyme that enters the secretory pathway and is delivered from the ER to the PPM. At the PPM, pro-PMII-GFP accumulates in cytostomes and migrates via vesicles to the DV.

**Figure 7:**
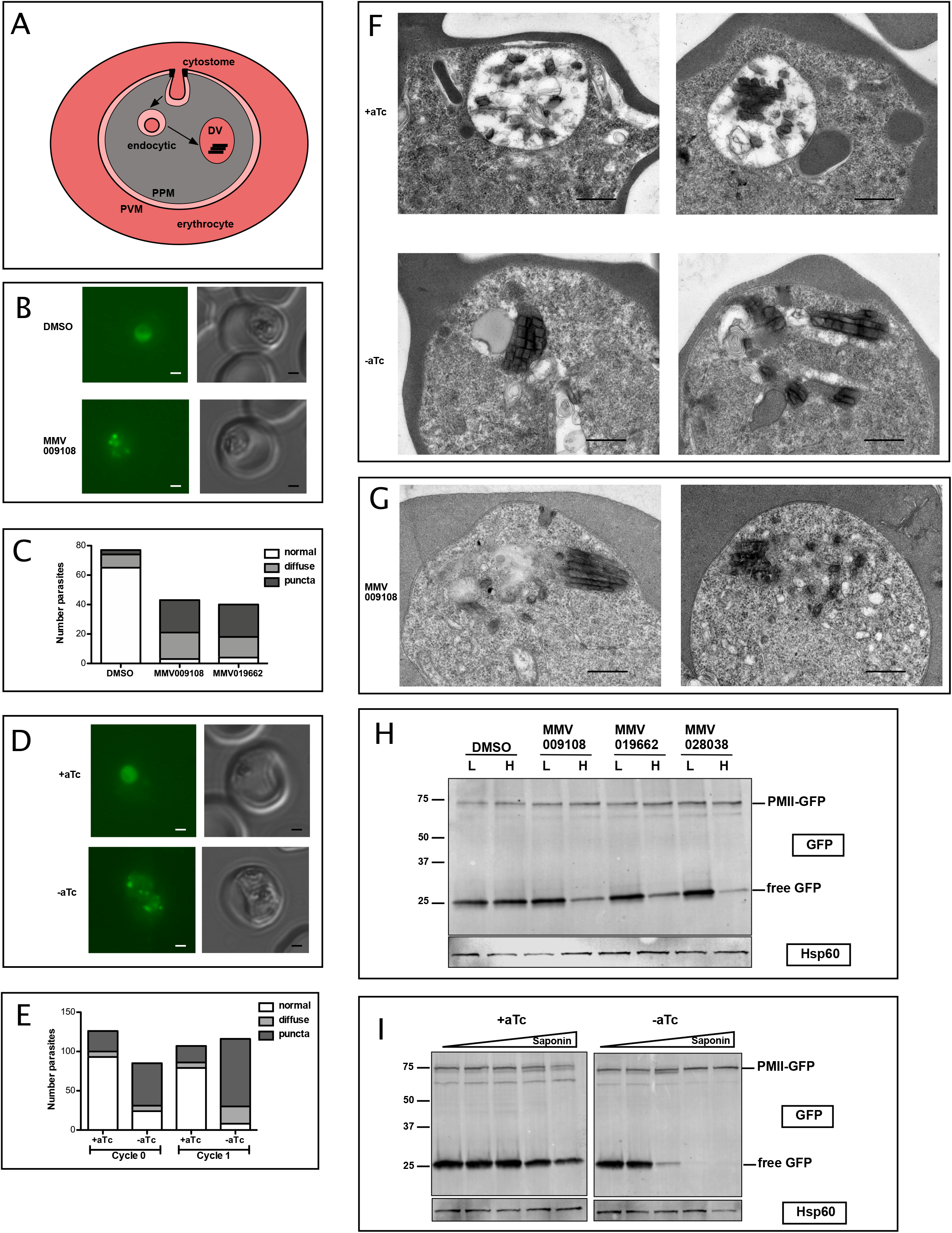
PfNCR1 inhibition or knockdown impairs digestive vacuole genesis. A) Cartoon of trafficking route to the DV in an infected red blood cell. DV = digestive vacuole; PVM = parasitophorous vacuolar membrane; PPM = parasite plasma membrane. B) Live microscopy of PMII-GFP parasites after incubation with MMV009108 (1μM, 3 hrs) or with vehicle (DMSO). Scale = 1μm. C) Quantitation of abnormal DVs from parasites in (B) after incubation with MMV009108, or MMV019662 or vehicle (DMSO) (1μM, 3 hrs). P<0.0001, Fisher’s exact test. D) Live microscopy of PfNCR1 K/D parasites expressing PMII-GFP, after removal of aTc. Scale = 1μm. E) Quantitation of abnormal DVs from parasites in (D) after aTc washout. P<0.0001, Fisher’s exact test. D and E): cycle 0 = trophozoites after removal of aTc within the same replication cycle (27hrs post washout), cycle 1 = trophozoites after removal of aTc in the preceding replication cycle (68 hrs post washout). F) Transmission electron micrographs of PfNCR1 K/D parasites (clone 2) after aTc removal (68 hrs post washout). Scale = 0.5μm. G) Transmission electron micrographs of PfNCR1 K/D parasites maintained with aTc and incubated with 500nM MMV009180 for 1 hr. Scale = 0.5μm. H) Western blot analysis of PMII-GFP parasites after treatment with 1μM compounds for 2 hrs. Parasites were released from RBCs with low (L) (0.009%) or high (H) (0.035%) saponin. Top blot was probed with α-GFP antibody, bottom blot (loading control) was probed with α-Hsp60 antibody, an organellar marker. This experiment was done two times. I) Western blot analysis of PMII-GFP, PfNCR1 K/D parasites after aTc washout for 22 hrs. Parasites were released from RBCs with 0.009%, 0.0175%, 0.035%, 0.07% or 0.14% saponin. Top blot was probed with α-GFP antibody, bottom blot (loading control) was probed with α-Hsp60 antibody. This experiment was done two times. Expected size of pro-PMII-GFP=79kDa, free GFP=27kDa.

After incubation with compounds we noticed abnormally punctate and occasionally diffuse GFP fluorescence that was not concentrated in DVs (Fig. 7B, C). Whereas most DMSO-treated control parasites had round DVs of ~2μm diameter and contained only a few small submicron GFP-positive dots, compound-treated parasites frequently had many small fluorescent foci, some of which were unusually bright. To confirm that abnormal DVs were a consequence of interfering with normal PfNCR1 function, we introduced PfNCR1^apt^ into the parasite line containing the PMII-GFP fusion (Supplementary Fig. S7). PfNCR1 K/D parasites had dispersed GFP puncta similar to those seen in compound-treated parasites (Fig. 7D, E). Electron micrographs prepared from parasites under PfNCR1 K/D (Fig. 7F) or treated with MMV009108 (Fig. 7G) showed dramatic defects. Normal DVs are easily distinguished from the parasite cytosol, not only because they contain hemozoin crystals, but also because they are electron-lucent. In contrast, the abnormal DVs we observed were electron-dense, smaller, elongated and irregular in shape. Usually, we could see multiple hemozoin-containing vesicles in PfNCR1-depleted/inhibited parasites.

To investigate whether DV membranes might contain defects similar to those observed in the PPM after PfNCR1 K/D or compound treatment, we measured the saponin sensitivity of the DV membrane. In PMII-GFP parasites, free GFP is hydrolyzed from PMII-GFP in the DV (Fig. 7A and ref.^25^). DV-resident GFP was released from drug-treated parasites at low saponin concentrations that did not affect control parasite DVs (Fig. 7H, I). Importantly, the levels of pro-PMII-GFP did not change, suggesting that the synthesis of PMII was not affected. To control for the possibility that DV membranes have increased leakiness after compound treatment or PfNCR1 K/D simply because the PPM is leaky and less detergent is necessary to access the DV, we repeated the experiment with isolated DVs. Again, after incubation with MMV009108, low saponin concentrations resulted in leakage of DV-localized GFP (Supplementary Fig. S7D). Metabolomic profiling of parasite extracts after incubation with MMV009108, MMV091662 (published previously^26^) or MMV028038 (Fig. 8) showed reductions across hemoglobin-derived peptides, supporting the hypothesis that the normal function of the DV has been compromised by the compounds.

**Figure 8:**
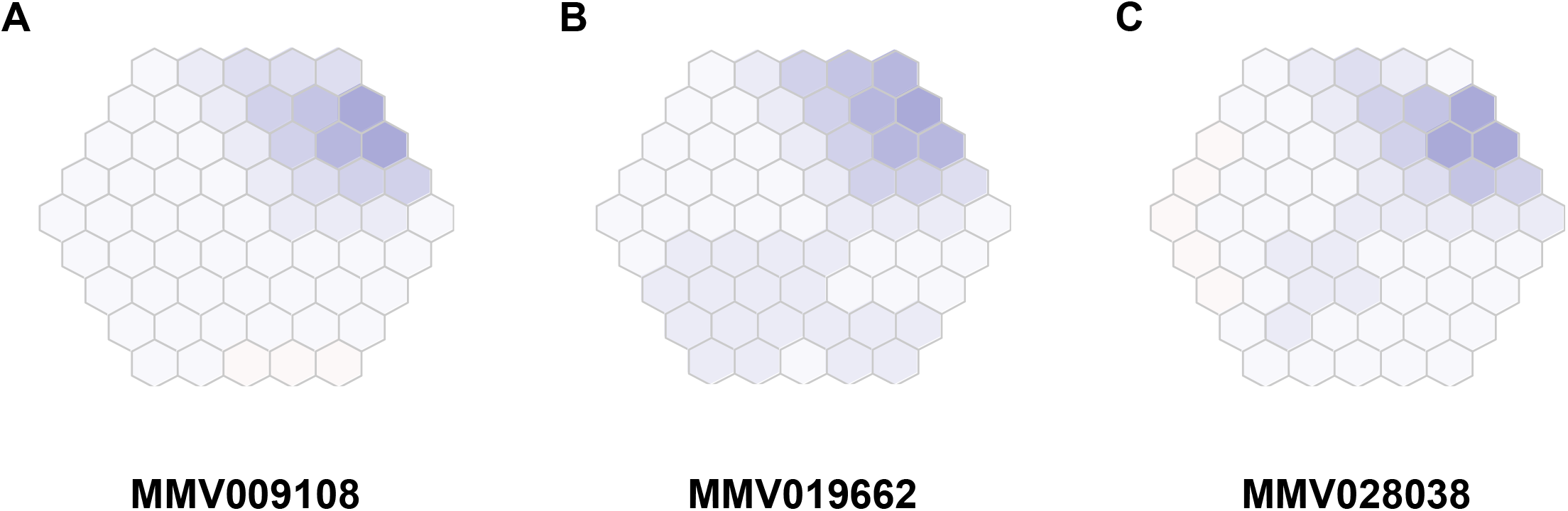
Metabolomic analysis of parasites incubated with PfNCR1 inhibitors. A)-C) Mass spectrometry-based metabolic profiling of hydrophilic extracts from parasites^26^ exposed to the three PfNCR1-targeting MMV compounds depicts a depletion in hemoglobin-derived peptides. Each panel represents incubation with a different compound. A) MMV009108, B) MMV019662, C) MMV028038. These Metaprint representations^48^ also demonstrate a highly similar metabolic response upon drug treatment with these compounds.

## Discussion

We have identified PfNCR1, Niemann-Pick C-related protein 1, as a new antimalarial target that resides in the PPM and serves essential functions during intraerythrocytic growth of *P. falciparum*. Through a chemical genetics approach we have provided evidence suggesting that three structurally diverse small molecules target PfNCR1. Conditional knockdown of *pfncr1* gene expression resulted in parasite demise. Phenotypic consequences of compound treatment or of conditional knockdown of PfNCR1 were essentially identical, strongly suggesting that the compounds directly inhibit PfNCR1.

PfNCR1 belongs to a superfamily of multi-pass transmembrane proteins involved in a variety of biological functions ranging from being receptors for signaling molecules to transport of different types of hydrophobic molecules^9,27,28^. Currently, the gene encoding this protein, PF3D7_0107500, is annotated as a lipid/sterol:H^+^ symporter (www.plasmodb.org). However, on the basis of its sequence similarity with previously investigated proteins from *Saccharomyces cerevisiae* (ScNCR1)^9^ and *Toxoplasma gondii* (TgNCR1)^14^ it is more appropriate to name it as PfNCR1. When engineered to display endosomal retention signals, ScNCR1 and TgNCR1 were able to revert defective cholesterol transport in mammalian cells lacking functional NPC1^14,29^. PfNCR1 displays 30% amino acid sequence identity over 69% of TgNCR1. Yet, there appear to be significant differences as to functions served by the proteins. Whereas ScNCR1 and TgNCR1 are dispensable for survival, PfNCR1 is essential. ScNCR1 has been localized to the yeast vacuole and *T. gondii* NCR1 to the inner membrane complex, a continuous patchwork of flattened vesicular cisternae located beneath the plasma membrane and overlying the cytoskeletal network; PfNCR1 is on the PPM.

Striking phenotypic consequences of PfNCR1 depletion or inhibition provide hints as to the functions served by this transmembrane protein. The ability of the cholesterol-dependent glycoside saponin to release cytosolic content of parasite-infected erythrocytes by permeation of the host plasma membrane while largely sparing the parasite cytosolic content has been a mainstay for experiments requiring “freeing” of parasites for biochemical and physiological investigations^30^. Cholesterol is not synthesized by malaria parasites but is taken up from the erythrocyte and incorporated into parasite membranes. An inward cholesterol gradient is formed as the parasite grows^31^. Resistance of the PPM to saponin permeation is believed to be due to a dearth of cholesterol within the PPM. Furthermore, the accessibility of cholesterol to saponin is highly dependent on its interactions with other lipids^32,33^. Interestingly, treatment with PfNCR1-active compounds results in saponin sensitivity of the parasites leading to the release of parasite cytosolic content within a short period of exposure. Remarkably, this saponin sensitivity was reversed upon the removal of the compounds targeting PfNCR1. The reversible saponin-sensitivity seen here is reminiscent of effects we have previously reported for antimalarial drugs that inhibit PfATP4, a P-type Na^+^ pump^22^. Induction of saponin sensitivity by PfATP4-active drugs was dependent upon the parasite being grown in a medium with standard [Na^+^]; saponin sensitivity was not seen in parasites grown in a medium with low [Na^+^]. Comparing the effects of PfNCR1-active compounds with PfATP4-active compounds, some similarities as well as differences become apparent. Both sets of compounds cause rapid but reversible saponin sensitivity in the PPM. PfATP4-active compounds disrupt Na^+^ homeostasis, which is a prerequisite for induction of saponin sensitivity, whereas PfNCR1-active compounds induce saponin sensitivity without disrupting Na^+^ homeostasis (Fig. 4F and Supplementary Fig. 5). It is possible that PfATP4 blockade perturbs the ionic environment critical for PfNCR1 function.

We noted that the concentrations at which PfNCR1-active compounds caused saponin sensitivity after a short exposure were much lower than the concentrations at which the compounds inhibited parasite growth in 72 h assays. Similarly, PfNCR1 knockdown caused saponin sensitivity of the PPM much sooner than inhibition of parasite growth. These results are opposite of what was previously seen for PfATP4-active compounds^22^. Parasites might have a greater tolerance for PPM composition disruption compared to the perturbation of Na^+^ homeostasis.

Another major consequence of PfNCR1 inhibition or knockdown was dramatic changes in formation and morphology of the DVs of parasites. DVs are lysosome-like organelles crucial for degrading hemoglobin. Unlike other eukaryotes or related apicomplexans, malaria parasites must actively digest hemoglobin to create room in the erythrocyte for the growing cell and to generate amino acids for parasite protein synthesis^34–36^. Uptake of erythrocyte cytosolic contents proceeds via the invagination of the PVM and the PPM and fusion of the PPM with the DV membrane contributes to mature DV formation^25^. Perhaps the abnormal membrane curvature^37^ and lack of fusion of the DVs upon loss/inhibition of PfNCR1 function provide clues towards understanding the critical requirement for normal lipid homeostasis in malaria parasites. The accumulation of hemoglobin peptides after incubation with PfNCR1 inhibitors suggests hemoglobin catabolism as a target pathway for these compounds and supports our findings. Among eukaryotes with NCR1 proteins but lacking receptor-mediated sterol uptake, malaria parasites are unusual in their requirement for functional NCR1 thus making this protein an exciting new antimalarial target. The diversity of chemical scaffolds targeting a single essential protein should provide guidance for future drug design and discovery efforts.

## Methods

### Parasite strains, culturing and resistance selection

Parasites were cultured in human red blood cells (2% hematocrit) in RPMI 1640 with 0.25% (w/v) Albumax (cRPMI) as previously described^25,38^. A lab-adapted strain of 3D7 that has been fully sequenced was used for most experiments^12^. For GFP overexpression in wild-type parasites, the previously described NF54^eGFP^ line was used, which bears an eGFP expression cassette targeted to the *cg6* locus using the attB x attP site-specific integrase recombination system^23^. Parasites with evolved resistance to MMV009108, MMV028038, or MMV019662 have been described^12^. Briefly, 5×10^8^ to 2×10^9^ 3D7 parasites were pressured with concentrations of 3x-10x EC_50_. Resistant parasites were readily obtained in multiple selections for the three compounds. Resistant and transfected parasites were cloned by limiting dilution. Dose-response experiments were done in triplicate starting with synchronous, young ring-stage cultures (1-1.2% starting parasitemia). Parasitemia (percentage of total erythrocytes infected with parasites) was measured approximately 70-80 hours post compound addition by nucleic acid staining of iRBCs with 0.8 μg/ml acridine orange in PBS. Growth was normalized to parasite cultures with carrier only (DMSO). Chloroquine (500nM) was used as a positive control for parasite growth inhibition. Data were fit to a sigmoidal growth inhibition curve. Growth curves of K/D and complemented parasites were done in biological triplicate with synchronous parasite cultures (aTc washout at young ring stage) by measuring daily parasitemias. Data were fit to an exponential growth equation. GraphPad Prism 5.0 was used for data analysis. Experiments for monitoring leakage (western blots and flow cytometry) of cytoplasmic HAD1, GFP or mRuby after compound treatment or under PfNCR1 K/D were performed with MACs LD (Miltenyi Biotech, Cat. No. 130-042-901) column-enriched parasites. Parasites were kept in cRPMI during all experiments. For aTc washouts, synchronous young ring-stage parasites were used. Washouts were repeated 3-4x, resuspending parasites at 2% hematocrit in cRPMI with 10min incubations at room temperature for each washout.

### Saponin release experiments

To monitor sensitivity to saponin, parasite cultures were pelleted (3 min x 840 g), pellets were suspended in 10X volume (most experiments) of room temperate saponin (prepared in PBS) for 2 mins (Sigma, Cat. No. S7900). Typical saponin concentration was 0.035%; modifications are indicated in the figure legends of experiments where appropriate. The released parasites were collected by centrifugation (3 min x 2200 g) and washed one time in cold PBS. In the experiment in Supplementary Fig. S4B (in which both supernatant and pellet fractions were collected) 2X volume saponin was used.

### Tetanolysin release experiments

Magnet-purified synchronous trophozoite-stage parasites were suspended in 10X volume of tetanolysin (0, 0.5, 1, 2.5, 5, 7.5ng/ml prepared in PBS) and incubated at room temperature for 2 min. The released parasites were collected by centrifugation (3 min x 2200 g) and washed one time in cold PBS.

### Cloning and Southern Blots

All plasmids were verified by direct Sanger sequencing.

#### 1) Allelic exchange constructs

Allelic exchange constructs were based on the vector pPM2GT^25^. Basepairs 2305-4893 of PF3D7_0107500 were cloned into the AvrII/XhoI sites using primers AR1-F and AR1-R primers. Using this strategy, *pfncr1* is expressed from the endogenous promoter in-frame with a C-terminal GFP (the native stop is deleted). The mutant constructs were prepared using QuikChange mutagenesis (Agilent Technologies, Cat. No. 20053). For the A1108T mutation, primer Mut-1 was used. In addition to the resistance mutation, this primer also introduces a BspHI site at bp 3294. For the A1208E mutation, primer Mut-2 was used. In addition to the resistance mutation, this primer also introduces a EcoRI site at bp 3605. For the F1436I mutation, primer Mut-3 was used. 100μg of circular DNA was transfected by electroporation of ring-stage parasites. Parasites were selected with 5nM WR99210 (kind gift of D. Jacobus), cycled twice off drug to enrich for parasites with integrated plasmid and cloned by limiting dilution.

#### 2) PfNCR1^apt^ parasites

In-Fusion cloning (Clontech) following PCR from gDNA was used to clone right and left homologous regions (RHR and LHR) for integration into the *pfncr1* locus. For the right homologous region, the sequence between bp3671 and bp4893 (the stop was deleted) (primers RHR1F and RHR1R) was amplified. An AflII site was introduced at the 5’ end and AatII was introduced at the 3’ end. Silent shield mutations to protect the construct from cleavage by CRISPR/Cas9 were introduced at S1464-S1465. For the left homologous region, a 948bp fragment starting 38bp past the stop codon was amplified (LHR1F and LHR1R). An AscI site was introduced at the 5’ end and an AflII site was introduced at the 3’ end. After generation of single homologous region fragments, RHR and LHR PCR products were mixed, amplified with primers RHR1F and LHR1R and cloned into the plasmid pMG75 as described^16^. The resulting construct (pMG75-PfNCR1) contains a single in frame HA sequence followed by 10x aptamers for aTc-regulatable translational repression. The construct contains two additional amino acids (D,V) before the HA sequence, as two tandem AatII sites were mistakenly introduced. For the gRNA sequence, the sequence 5’-TTAATGTAGTGGGCCAAAAC-3’ was chosen. The sense and antisense primer pair GRNA1 and GRNA2 encoding the *pfncr1* sgRNA seed sequence was annealed and inserted into the BtgZI site in plasmid pyAIO^16^, resulting in the plasmid pyAIO-PfNCR1-gRNA1. 100μg of pMG75-PfNCR1 was linearized with AflII, purified by phenol-chloroform extraction and co-transfected with 50μg of pyAIO-PfNCR1-gRNA1 by electroporation. Parasites containing the modified *pfncr1* locus were selected with 5μg/ml Blasticidin S. For the PfNCR1^apt^ strain that expresses PMII-GFP, we transfected the previously described PMII-GFP clone^25^ with pyAIO-PfNCR1-gRNA1 and linearized pMG75-PfNCR1. In this case, parasites were selected with 5nM WR99210 plus 5μg/ml Blasticidin S and kept in media with 500nM aTc. Parasites were cloned by limiting dilution.

#### 3) Complementation of PfNCR1^apt^

For complementation, RNA was prepared from 3D7 parasites using TRIzol (ThermoFisher), *pfncr1* RNA was amplified using a SuperScript RT-PCR kit (Invitrogen) with primers Comp1 and Comp2, cloned into the XhoI/AvrII sites of the pTEOE random integration vector with the PiggyBac transposase as described^39,40^. PfNCR1^apt^ clone 2 was transfected and selected with 5μg/ml Blasticidin S and 2μM DSM-1 (Asinex)^41^.

#### 4) Expression of cytoplasmic GFP in PfNCR1^apt^ background

GFP overexpression in PfNCR1^apt^ parasites was achieved by targeting the eGFP expression cassette of NF54^eGFP^ parasites to the *rh3* locus by CRISPR/Cas9 editing. The *calmodulin* promoter and *egfp* coding sequencing was amplified from NF54^eGFP^ genomic DNA template using primers eGFP-F and eGFP-R and inserted into the plasmid pPM2GT^25^ between AatII and EagI by In-Fusion cloning, allowing for fusion to the *hsp86* 3’ UTR. The sgRNA target site TGGTAATACAGAAATGGATG was chosen in the dispensable *rh3* gene. Homology flanks were then amplified from sequence just upstream and downstream of the Cas9 cleavage site defined by this sgRNA using primers Rh3-5’F/R and Rh3-3’F/R. These amplified flanks were used as template and assembled into a single DNA molecule with an intervening AflII site in a second PCR reaction using primers Rh3-5’R and Rh3-3’F and this flank assembly was inserted into the BglII site of the pPM2GT-CAM-eGFP plasmid resulting in the plasmid pPM2GT-CAM-eGFP-RH3-flanks. A sense and antisense primer pair (Rh3-G1 and Rh3-G2) encoding the *rh3* sgRNA seed sequence was annealed and inserted into the BtgZI site in plasmid pyAIO^16^ resulting in the plasmid pyAIO-RH3-gRNA1. Plasmid pPM2GT-CAM-eGFP-RH3-flanks was linearized at AflII and co-transfected with pyAIO-RH3-gRNA1 into PfNCR1^apt^ clone 2 and selected with 2μM DSM-1 for integration into the rh3 locus.

#### 5) Expression of split-GFP

For split GFP experiments, two parasite lines were generated expressing either PV-targeted or cytosolic GFP1-10. A fusion of the *sera5* signal peptide and *gfp1-10* coding sequence was synthesized as a gBlock (gBlock1; IDT) and used as template to PCR amplify *gfp1-10* with primers GFP1-10-1F and GFP1-10-1R or without the *sera5* signal peptide (primers GFP1-10-2F and GFP1-10-1R). These amplicons were inserted into plasmid pLN-ENR-GFP^42^ between AvrII and AflII to generate plasmids pLN-SP-GFP1-10 and pLN-GFP1-10, respectively. Each plasmid was co-transfected with plasmid pINT into NF54^attB^ parasites and selected with 2.5 μg/ml Blasticidin S to facilitate integration into the *cg6* locus through integrase-mediated attB x attP recombination^42^. A clonal line was derived from each transfected parasite population by limiting dilution and designated NF54^pvGFP1-10^ or NF54^cytGFP1-10^, respectively. GFP1-10 expression and targeting to the proper compartment (parasitophorous vacuole or cytosol) was confirmed by western blot and immunofluorescence assay using a rabbit-anti-GFP (Abcam 6556). For endogenous tagging of PfNCR1 with 3xHA-GFP11, *pfncr1* was amplified from pMG75-PfNCR1 with primers GFP11-F and GFP11-R and inserted into the plasmid pyPM2GT-EXP2-mNeonGreen^43^ between XhoI and AvrII. Transfections were selected with 2μM DSM-1. This construct expresses PfNCR1-GFP11 from its native promoter.

#### 6) For monitoring PVM integrity

The line NF54-EXP2-mNeonGreen+PV-mRuby3 was used^44^.

### Southern blot

To confirm correct integration, we used the AlkPhos Direct Kit (FisherScientific Cat. No. 45-000-936) for Southern blots as described^25^. For the probe, we amplified a 674 bp fragment from gDNA using primers Probe1 and Probe2.

### Microscopy

Fluorescence microscopy was performed on live, GFP-expressing parasites using a Zeiss Axioskope. Nucleic acid was detected by staining with DAPI.

For electron microscopy, infected RBCs were enriched using MACs LD columns, fixed in 4% paraformaldehyde (Polysciences Inc., Warrington, PA) in 100mM PIPES/0.5mM MgCl_2_, pH 7.2 for 1 hr at 4°C. Samples were then embedded in 10% gelatin and infiltrated overnight with 2.3M sucrose/20% polyvinyl pyrrolidone in PIPES/MgCl_2_ at 4°C. Samples were trimmed, frozen in liquid nitrogen, and sectioned with a Leica Ultracut UCT cryo-ultramicrotome (Leica Microsystems Inc., Bannockburn, IL). 50 nm sections were blocked with 5% FBS/5% NGS for 30 min and subsequently incubated with rabbit anti-GFP (Life Technologies; Cat. No. A11122) (1:500) for 1 hr, followed by goat anti-rabbit IgG (H+L) antibody conjugated to 18 nm colloidal gold (1:30) (Jackson ImmunoResearch) for 1 hr. Sections were washed in PIPES buffer followed by a water rinse, and stained with 0.3% uranyl acetate/2% methyl cellulose and viewed on a JEOL 1200EX transmission electron microscope (JEOL USA, Peabody, MA) equipped with an AMT 8 megapixel digital camera (Advanced Microscopy Techniques, Woburn, MA). All labeling experiments were conducted in parallel with controls omitting the primary antibody which was consistently negative at the concentration of colloidal gold conjugated secondary antibodies used in these studies. For EM without immunostaining, cells were fixed in 2% paraformaldehyde/2.5% glutaraldehyde (Polysciences Inc., Warrington, PA) in 100 mM sodium cacodylate buffer, pH 7.2 for 1 hr at room temperature. Samples were washed in sodium cacodylate buffer and postfixed in 1% osmium tetroxide (Polysciences Inc.) for 1 hr. Samples were then rinsed extensively in dH_2_O prior to en bloc staining with 1% aqueous uranyl acetate (Ted Pella Inc., Redding, CA) for 1 hr. Following several rinses in dH_2_O, samples were dehydrated in a graded series of ethanol and embedded in Eponate 12 resin (Ted Pella Inc.). Sections of 95 nm were cut with a Leica Ultracut UCT ultramicrotome (Leica Microsystems Inc., Bannockburn, IL), stained with uranyl acetate and lead citrate, and viewed on a JEOL 1200 EX transmission electron microscope (JEOL USA Inc., Peabody, MA) equipped with an AMT 8 megapixel digital camera and AMT Image Capture Engine V602 software (Advanced Microscopy Techniques, Woburn, MA).

### Flow Cytometry

For flow cytometry experiments with eGFP, 50,000 cells were counted on a BD FACSCanto and scored for high or low GFP signal. Appropriate gating of cells was established using untreated parental or NF54^eGFP^ parasites.

### Western blotting

For PfNCR1 blots, membrane preparations were made. 1×10^8^ to 5×10^8^ trophozoite-stage parasites were released from RBC with 0.035% saponin, washed in cold PBS, resuspended in 300μl DI-water with protease inhibitors (HALT, ThermoFisher, Cat. No. 78430), freeze-thawed 3x with liquid nitrogen/42^0^C water bath. The membranes were pelleted (17k g), resuspended in 100μl-300μl (depending on sample amount) Ripa buffer (25mM Tris (pH 7.6), 150mM NaCl, 1% NP-50, 0.1% SDS, 1% Sodium Deoxycholate) containing 0.1% CHAPS and 0.1% ASB-14, sonicated 3x with a microtip, and incubated at 42^0^C with shaking for 45min. The samples were then centrifuged (17k g, 30min), SDS sample buffer was added to the soluble portions. The samples were warmed at 42^0^C and loaded on 4-15% TGX gradient gels (Biorad). Proteins were transferred onto PVDF using wet transfer with 20% methanol. Blots were blocked either 1hr at 25^0^C or overnight at 4^0^C with Licor Odessey block buffer. Primary antibodies were mouse monoclonal α-HA antibody (Biolegend) at 1:1000 or LivingColors mouse-α-GFP (Takara, Cat. No. 632380) (1:1000). For the loading control mouse monoclonal α-PM-V antibody^45^ at 1:20 was used. Secondary antibody was goat-α-mouse (800) IR-Dyes (1:20,000) from Licor.

For western blot monitoring leakage of cytosolic proteins after incubation with compound or PfNCR1 K/D, parasites were resuspended in saponin-containing PBS, pelleted, lysed in Ripa buffer containing protease inhibitors and with brief sonication. Soluble proteins after centrifugation (30min, 17k g) were added to sample buffer, briefly heated at 98^0^C and loaded onto 10% or 12% TGX gels (Biorad). Western blotting was done using the protocol indicated above. Primary antibodies were: rabbit α-HAD1 (a gift from Dr. Audrey Odom John, WU)^46^ (1:1000), rabbit α-Hsp60 (1:500) (a gift from Dr. Sabine Rospert, University of Freiburg), mouse α-PM-V (1:20)^45^, rabbit α-RFP (1:1000) (Thermofisher, Cat. No. R10367), mouse α-GFP (Living Colors JL-8, Clontech, Cat. No. 632380) (1:1000), HRP-conjugated α-aldolase (Abcam, Cat. No. ab38905) (1:10000), α-EXP2 antibody (gift from Professor James Burns, Drexel University) (1:10000)^22^. Secondary antibodies were goat-α-mouse (800) and donkey-α-rabbit (680) IR-Dyes (1:20,000) from Licor. Immunoblots shown in Figs. 4A, 4E and Supplementary Fig. S6 were washed in PBS-Tween (0.2%) and developed using the Super Signal West Pico Chemiluminescent substrate (Thermo Scientific, Cat. No. 34080).

### Measuring intracellular [Na^+^]

Effect of compounds on intracellular [Na^+^] was determined using methods described by Spillman et al.^47^. Saponin freed parasites were loaded with the ratiometric sodium-sensitive probe SBFI-AM (5.5 μM) (Invitrogen) and 0.01% w/v Pluronic F-127 (Invitrogen) in suspension (at 2.5-3.5 x 10^8^ parasites/mL) for 30min at 37°C in bicarbonate-free RPMI supplemented with 20 mM glucose, 10 mg/L hypoxanthine, 25 mM HEPES and 50 mg/L gentamycin sulphate (pH 7.1). The probe-loaded parasites were washed twice (2,000x g, 30 s) and resuspended to a final parasite concentration of 1.0-1.5 x 10^8^/mL in a saline buffer (125 mM NaCl, 5 mM KCl, 1 mM MgCl_2_, 20 mM glucose, 25 mM HEPES, pH 7.1). SBFI loaded parasites were excited at 340 nm and 380 nm with emissions recorded at 500 nm in a Horiba Scientific FluoroMax 3 spectrofluorometer for 5 min followed by addition of the compounds at indicated concentrations. Auto-fluorescence corrected 340/380 nm emission fluorescence ratio were related to [Na^+^]_i_ using an average from 3 independent calibration curves for SBFI. Calibration plots were generated for SBFI-loaded parasites in solutions containing 0, 10, 25, 50, 75, 100, 130 mM Na^+^, made by mixing solutions of 80 mM Na^+^/K+ gluconate and 50 mM NaCl/KCl (1 mM MgCl_2_, 20 mM glucose, 25 mM HEPES, pH 7.1) and [Na^+^]_i_ was equilibrated using a combination of the ionophores nigericin (5 μM), gramicidin (2.5 μM) and monensin (5 μM).

### Metabolomic Profiling

Changes in metabolites were measured in response to compounds using whole cell hydrophilic extraction, followed by ultra-high precision liquid chromatography mass-spectrometry (UHPLC-MS) using negative ionization as in Cowell et al., 2018^13^. This was performed on synchronous, trophozoite infected red blood cells (iRBCs, 24-36 hpi) which had been magnetically separated from culture. Quantification of cells was performed by hemocytometry, and treatments were performed on 1×10^8^ iRBCs in wells containing 5 mL of RPMI. Treatment conditions were performed in triplicate, with compound concentrations of 10×EC_50_ for 2.5 hours, followed by washing with PBS and extraction using 90% methanol containing isotopically-labeled aspartate as an internal standard for sample volume. Samples were dried using nitrogen prior to resuspension in water containing 0.5 uM chlorpropamide as an internal standard for injection volume. Samples were then analyzed via UHPLC-MS on a Thermo Scientific EXACTIVE PLUS Orbitrap instrument as established in Allman et al., 2016^26^.

## Acknowlegdements

We are thankful to our MalDA Consortium collaborators and D. S. Ory (WU) for stimulating discussions, A. S. Nasamu and A. Polino for valuable suggestions, B. Vaupel for assistance during cloning, W. Beatty for electron microscopy, and M. Lee, M. Carrasquilla and J. Rayner for consulting on the Rh3 gRNA design.

This work was supported by Gates Foundation Grants OPP 1054480 (Winzeler, Goldberg, Llinás), OPP1132313 (Niles), and NIH grants R01AI098413 and R01AI132508 (Vaidya); K99/R00 HL133453 (Beck), and 1DP20D007124 (Niles).

